# A comparative study of metagenomics analysis pipelines at the species level

**DOI:** 10.1101/081141

**Authors:** Yee Voan Teo, Nicola Neretti

## Abstract

Many metagenomics classification tools have been developed with the rapid growth of the metagenomics field. However, the classification of closely related species remains a challenge for this field. Here, we compared MetaPhlAn2, kallisto and Kraken for their performances in two metagenomics settings, human metagenomics and environmental metagenomics. Our comparative study showed that kallisto demonstrated higher sensitivity than MetaPhlAn2 and Kraken and better quantification accuracy than Kraken at the species level. We also showed that classification tools that run on full reference genomes misidentified many species that were not truly present. In order to reduce false positives, we introduced marker genes from MetaPhlAn2 into our pipeline, which uses kallisto for the classification step, as an additional filtering step for species detection.

## Introduction

The advent of shotgun metagenomic sequencing greatly facilitated the identification and classification of microbes by providing a means to detect phenotypically aberrant or unculturable microbes [1]. It allows a much faster and cheaper taxonomic profiling of microbial communities in different ecosystems such as the microbiome in human, soil and ocean. With the rising use of shotgun metagenomic sequencing in the last decade, many microbial species classification tools have since been developed [2]. However, there still remains one of the main challenges in shotgun metagenomics analysis—the genomes similarity problem among closely related species in which it is hard to distinguish and classify ambiguous reads. [3]. For instance, *Shigella dysenteriae* and *Escherichia coli* that share relatively similar genomes complicate the taxonomic assignment at the genus, species and strain levels [4]. Several tools such as MetaPhlAn and MetaPhyler have been developed to profile microbial communities rapidly using a set of marker sequences. The use of markers can reduce ambiguous reads mapping to multiple genomes [5-8], but at the same time, not all sequencing reads can be classified. This poses a limit to perform a detailed analysis on the samples, such as gene content estimations [9]. Other tools such as Kraken [9]and Clark [10] also have been developed for high accuracy microbial sequence classification. These tools represent read-alignment free and k-mer based approach that can classify sequencing reads accurately and rapidly. Lindgreen et al. evaluated many of the widely used metagenomics classification tools and the comparison of the overall performances showed that Kraken performs best in terms of the speed and accuracy in identifying taxonomic distribution [11]. Interestingly, an RNA-seq quantification tool, kallisto, has also been tested and compared to Kraken in the metagenomics setting. Kallisto is a fast k-mer based pseudoalignment approach of RNA-seq reads to quantify isoform expression level using a transcriptome De Bruijn graph (T-DBG) method [12]. The comparison of Kraken and kallisto by Schaeffer et al. showed that kallisto outperforms Kraken in the metagenomics quantification at three taxonomic ranks: genus, species and strain [13]. However, no evaluations on other aspects such as the false discovery rates (FDR) and memory requirements have been reported in the study.

Here, we compared and evaluated different combinations of tools, including MetaPhlAn2, kallisto and Kraken, in terms of their detection and quantification performances, speed and memory requirement in microbial species level classification. We showed that the performance of kallisto is better than Kraken in terms of the quantification at the species level, which is consistent with the result showed by Schaeffer et al [13]. However, while kallisto uses expectation maximization (EM) algorithm to probabilistically handle ambiguous reads, the FDR still increases dramatically as the number of sequencing reads increases. Therefore, we incorporated a collection of species-specific markers genes from MetaPhlAn2 into the kallisto quantification pipeline as an additional step to reduce the FDR. In addition, due to the very high memory requirement to build the T-DBG by kallisto, we also introduced another pipeline that build the index only on detected species from the kallisto run on marker genes to evaluate the performance of kallisto at a larger scale of reference genomes. Overall, we showed that the kallisto run on full microbial genome alone is not sufficient because it detected many other species that were not there. The use of marker genes is necessary to reduce the high false positives.

## Results

Two different microbial community samples were simulated: (i) human-associated habitat microbial community samples that consist of 5% microbial reads, and (ii) samples that consist of only microbial reads. We tested five pipelines that use either MetaPhlan2, Kraken or kallisto (Figure 9 and Figure 10) for the classification of reads at the species level and evaluated their performances in terms of sensitivity, FDR, rate of false negative and memory requirement.

### (i) Human-associated habitat microbial community samples

For datasets that mimic samples extracted from human-associated habitat, we first filtered out human reads before running the classification step. Sensitivity measures how well the pipelines detect species that were truly present in the samples. Pipelines that include kallisto (Figure 9 B1-D1) in the classification step demonstrated the highest sensitivity, followed by Kraken (Figure 9 E1) and MetaPhlAn2 (Figure 9 A1, Figure 1a). MetaPhlAn2 performed poorly at low number of microbial reads due to the use of only marker genes and not full genomes. FDR measures the number of species that were detected but not in the samples. It increases dramatically as the number of reads increases when full microbial genomes were used in the reference database (Figure 1b). In contrast, pipelines that involve additional filtering step using species-specific marker genes from MetaPhlAn2 (C1 and D1) and MetaPhlAn2 itself (A1) demonstrated lower FDR (<0.05) compared to the pipelines that only classify reads using full microbial genomes. Rate of false negative refers to the frequency of not detecting species that were present in the samples. Pipelines that include kallisto (B1-D1) showed the lowest rate of false negative, followed by Kraken (E1) and MetaPhlAn2 (A1) (Figure 1c). Although kallisto performed well in the detection of species, the downside of it is that it requires high memory in the index and quantification steps. Specifically, kallisto quantification (B1 and C1) with a database that consists of 3511 fasta sequences (~5.6GB) consumed approximately 300 GB of memory (Figure 1d). Therefore, out of the five pipelines that we tested, we included one that index only genomes that were detected in the marker genes detection step (D1), in order to reduce the size of the database used in the final quantification. In this case, the memory requirement will vary depending on the number of species detected. The number of genomes in our simulated dataset was 315 and kallisto required approximately 38GB of memory for the quantification step. In addition, the run time also varies between different pipelines (Figure 2a-e). On a 16-core server, MetaPhlAn2 demonstrated the shortest run time due to its small database that consist of only marker genes. Kraken pipeline had the second shortest run time, followed by the three kallisto pipelines, with B1 and C1 that run on the full microbial genome having the longest run time. Similar to the memory requirement by the D1 pipeline, the run time that involves building a kallisto index and running on a reduced database will vary depending on the number of species detected.

**Figure 1:**
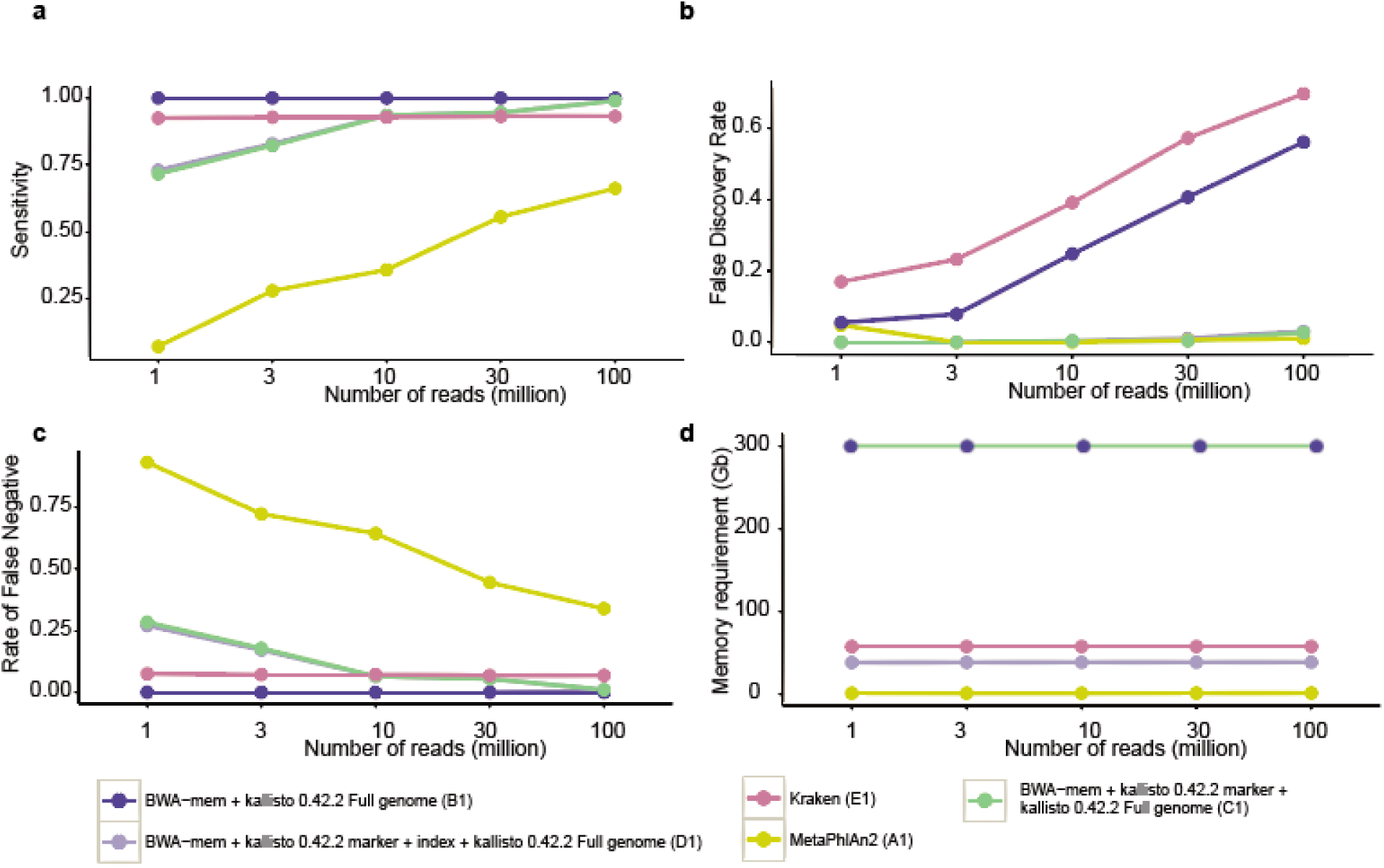
**(a)** Sensitivity, **(b)** false discovery rate, **(c)** rate of false negative and **(d)** memory requirement for simulated microbial reads extracted from human-associated habitat

**Figure 2.**
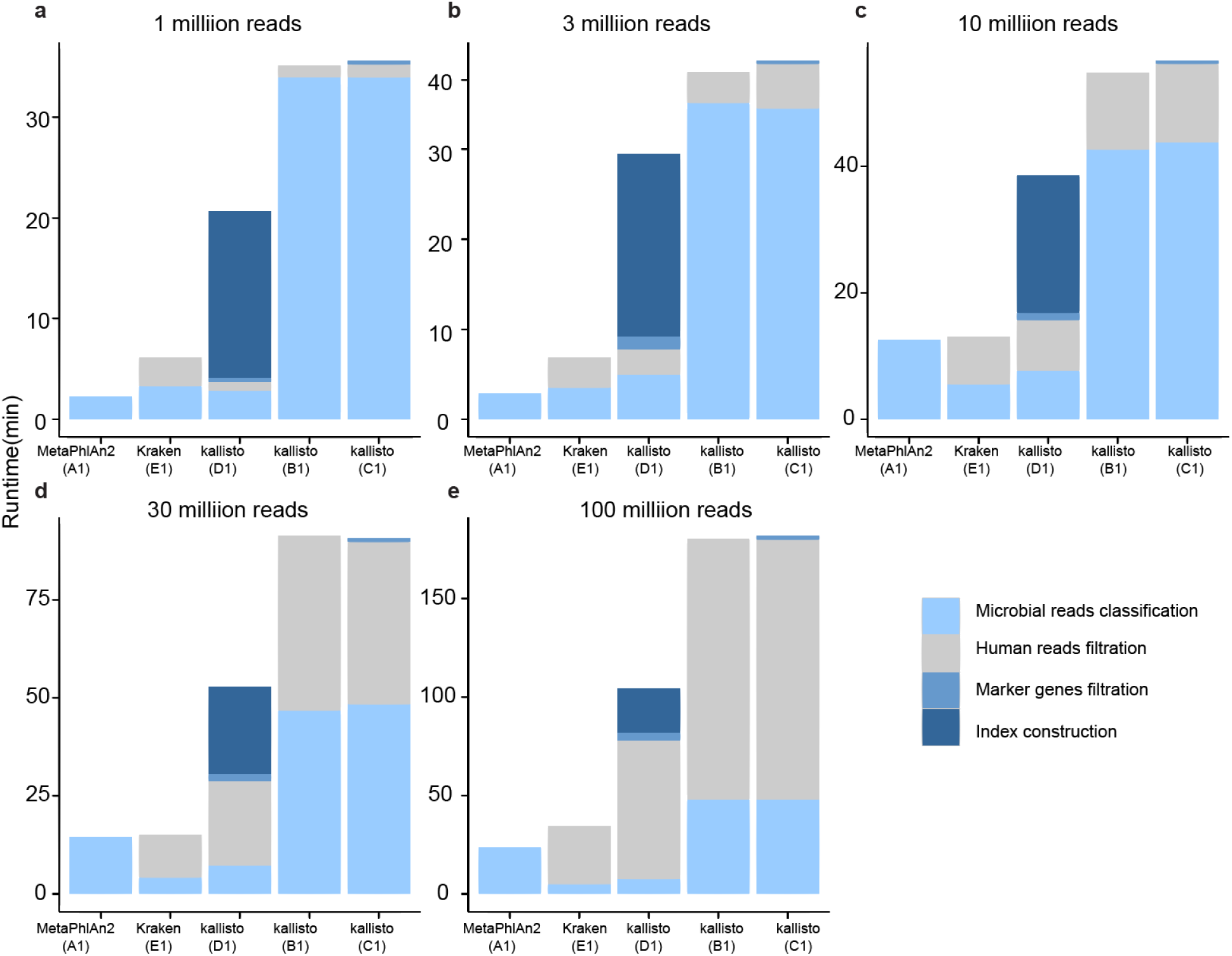
Runtime (min) for samples extracted from human-associated habitat that consist of **(a)** 1 million, **(b)** 3 million, **(c)** 10 million, **(d)** 30 million, and **(e)** 100 million total reads

We excluded MetaPhlAn2 from the comparison of quantification accuracy because not all reads are used in the estimation of microbial composition. The kallisto pipeline (D1) that has a filtration step with species-specific markers demonstrated the highest accuracy in estimating species counts as the number of reads increases, followed by the other two kallisto pipelines (C1 and B1), and then Kraken (E1) (Figure 5a). In summary, we showed that the D1 pipeline demonstrated the most optimal combination of all parameters tested although it takes a longer time to run compared to MetaPhlAn2 and Kraken. It first filters out human reads using BWA-MEM, followed by a species level detection step using marker genes, and a quantification step on a selected full genome database based on detected species.

### (ii) Samples of only microbial reads

The same five pipelines used in the first part of the study minus the human filtration step were evaluated for their performances in classifying samples with only microbial reads. The performances of each pipeline were similar to the first part of the study (Figure 3a-d), in which pipelines that include kallisto (Figure 10 B2-D2) in the classification step demonstrated the highest sensitivity, followed by Kraken (Figure 10 E2) and MetaPhlAn2 (Figure 10 A2). Pipelines that do not have a filtering step using species-specific marker genes (B2 and E2) demonstrated higher FDR as the number of reads increases. Kallisto pipelines (B2 and C2) are not practical because it requires memory as high as 300GB for a database of only 3511 microbial reference sequences (Figure 3d).

**Figure 3:**
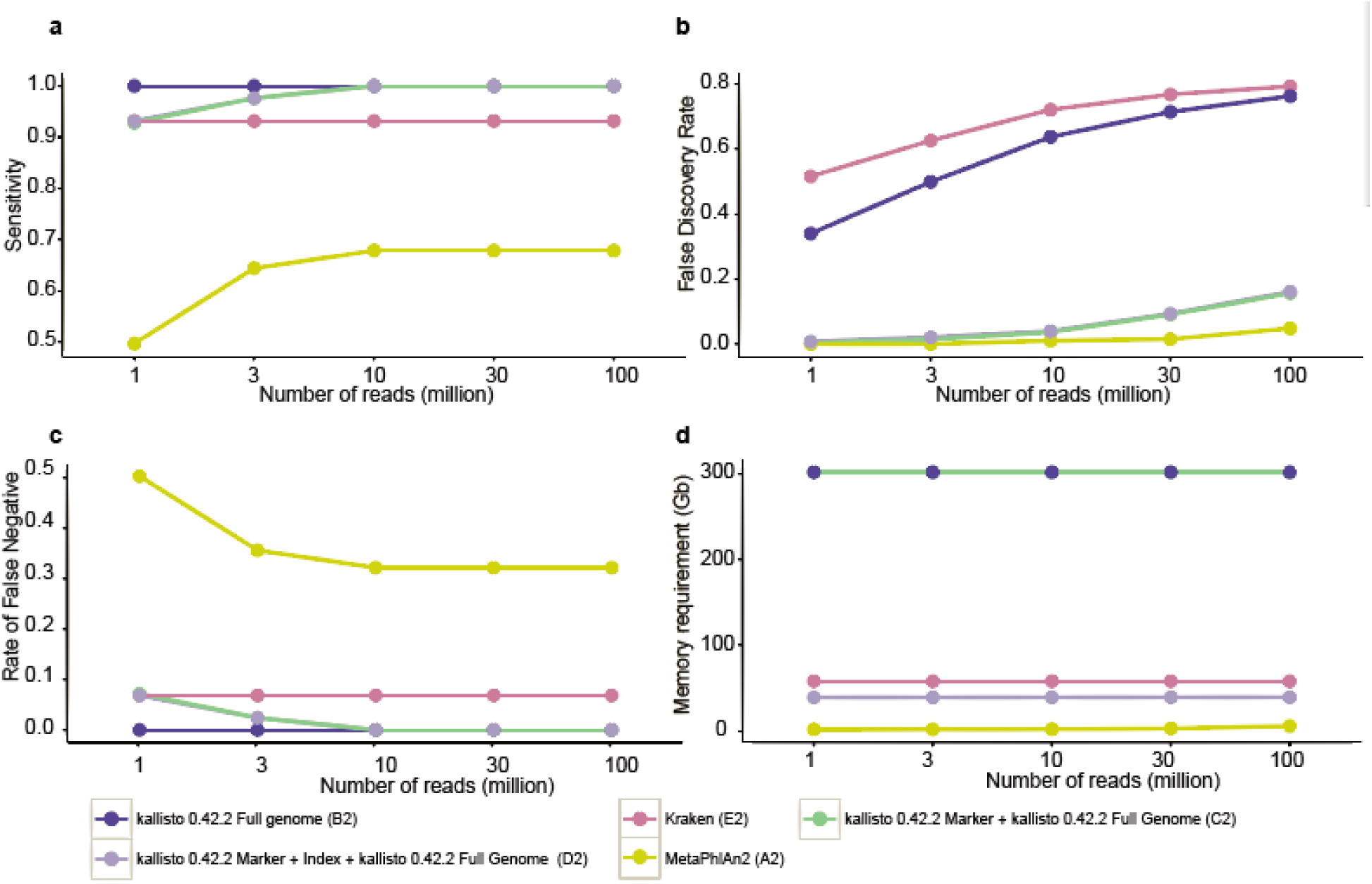
**(a)** Sensitivity, **(b)** false discovery rate, **(c)** rate of false negative and **(d)** memory requirement for simulated samples that contain only microbial reads

In terms of the speed, MetaPhlAn2 (A2) had the shortest run time on a 16-core server. However, as the number of reads increases, Kraken (E2) outperformed MetaPhlAn2 in terms of the run time (Figure 4a-e) even though it runs on a full reference genome database. Kraken could classify 100 million reads in about 30 minutes. Kallisto (B2 and C2) had the longest run time for microbial reads classification when full reference genomes database was used.

**Figure 4:**
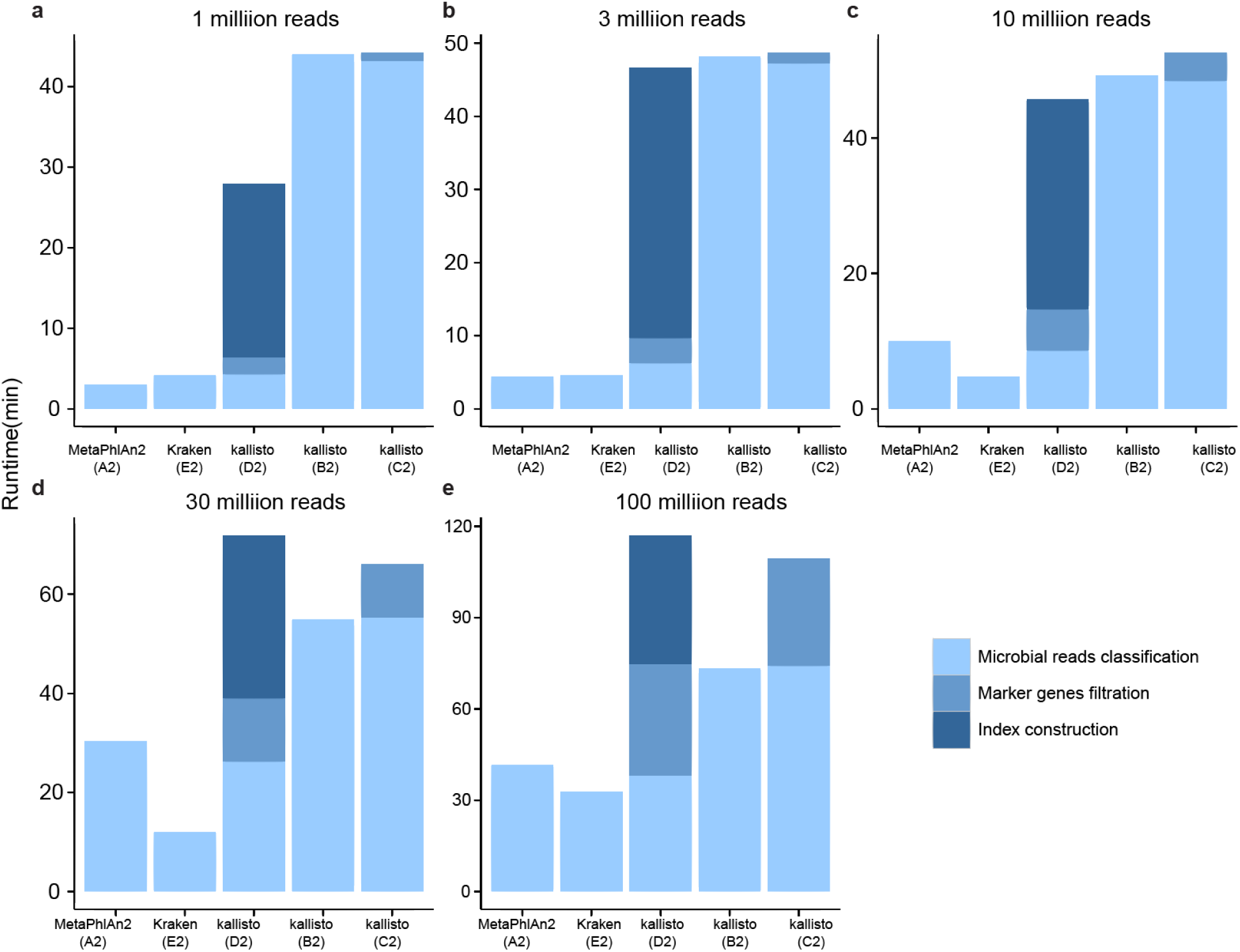
Runtime (min) for simulated samples that consist of **(a)** 1 million, **(b)** 3 million, **(c)** 10 million, **(d)** 30 million, and **(e)** 100 million microbial reads

**Figure 5:**
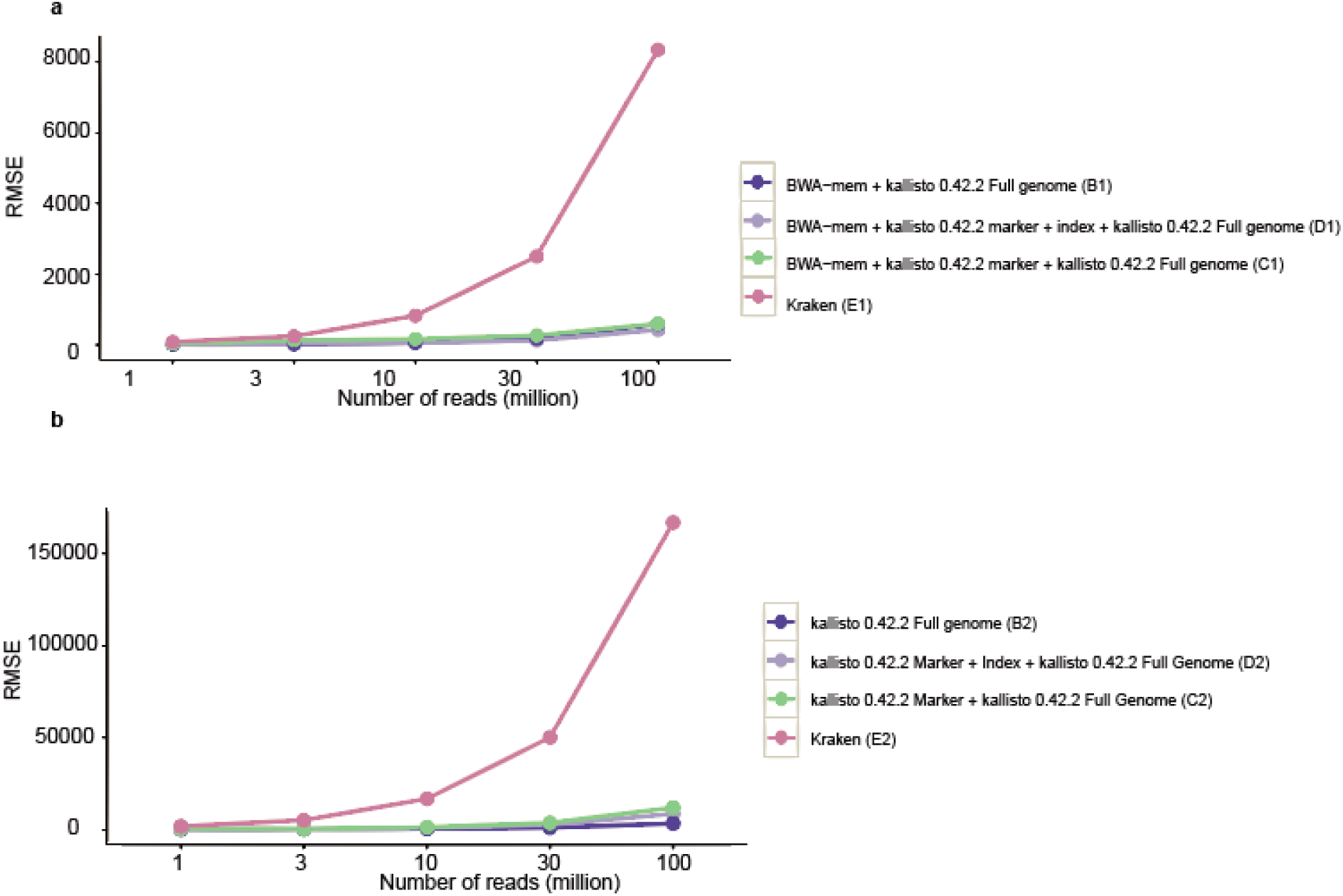
RMSE of 292 organisms (**a)** from human-associated habitat and **(b)** of all microbial reads only, at the species level

The kallisto pipeline (B2) without the filtration step with species-specific markers demonstrated the highest accuracy in estimating species counts, followed by the other two kallisto pipelines (D2 and C2), and then Kraken (E2). The downside of this B2 pipeline was that the FDR increases dramatically to as high as 0.76 with 100 million of microbial sequencing reads (Figure 3b). This indicates that kallisto quantification alone is not sufficient to accurately identify the species that are present in a sample and the use of marker genes is necessary to reduce the FDR. The additional marker genes filtration step will definitely require more run time than without one. The quantification step by kallisto also consumes a much longer time (10 times as long when quantifying 1 million microbial reads) when compared to Kraken (Figure 4a). Overall, in order to get an accurate quantification of microbial reads, yet the highest sensitivity and the lowest FDR, the D2 pipeline demonstrated the most optimal combination of all parameters tested, although the run time is not the best. The first step is the species level detection using marker genes to reduce the FDR, followed by a quantification step on a selected full genome database based on detected species.

### (iii) iMESS_Illumina simulated Illumina100

In addition to our simulated sequencing reads, we also tested the D2 pipeline on the dataset that was used by Schaeffer et al. to test kallisto in a metagenomics setting. 56.9% of the reads are from strains found in our microbial reference genomes database, 34.1% of them are of other strains that can be classified into species found in the database, and the remaining 9% are not in any of the species found in the database. We wanted to test how well the D2 pipeline can classify when some of the reads are not found in the database because this can mimic the problem in analyzing real dataset. The pipeline correctly identified 67 species out of the 85 species that were truly present in the dataset and misidentified 72 species. The quantification accuracy of this approach was shown in Figure 6 (Pearson r = 0.75).

**Figure 6:**
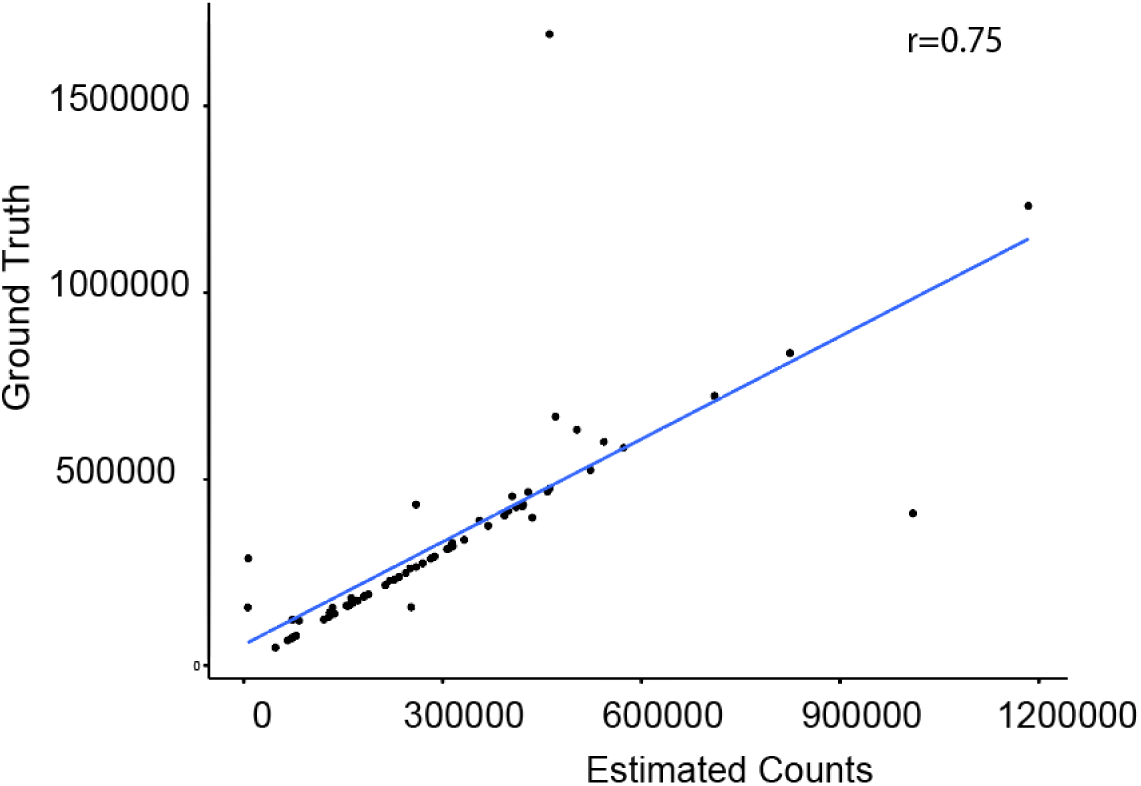
Quantification accuracy for Illumina100 dataset using the D2 pipeline

### (iv) Stool and saliva sequence data from the HPFS cohort

We obtained stool and saliva samples from eight healthy Health Professionals Follow-Up Study (HPFS) subjects and analyzed using the D1 pipeline. There was a clear clustering structure (Figure 7) between the oral and the gut microbiome of the eight HPFS subjects. The most distinctive species composition that distinguish between the oral and the gut microbiome were members from the *Prevotella* genus and the *Bacteroides* genus (Figure 8a). Members in the *Prevotella* genus such as *Prevotella histicola, Prevotella melaninogenica, Prevotella pallens* and *Prevotella sp. C561* were found in all eight oral samples but not in any of the gut sample (FDR < 3.6 x 10^−16^). In contrast, species in the *Bacteroides* genus such as *Bacteroides sp. 9 1 42FAA, Bacteroides sp. HPS0048, Bacteroides xylanisolvens, and Bacteroides nordii* were not found in the oral samples but were found in the gut samples (FDR < 3.1 x 10^−5^). In addition, we also identified two species, *Veilonella sp. HPA0037* and *Streptococcus salivarius* with at least 1000 counts per million (CPM), in both the gut and the oral samples from the same individual, in at least two of them. The link between the oral and the gut microbiome from the same dataset has previously been shown in [14] and our finding on *Streptococcus salivarius* and the *Veilonella* genus was consistent with their result.

**Figure 7:**
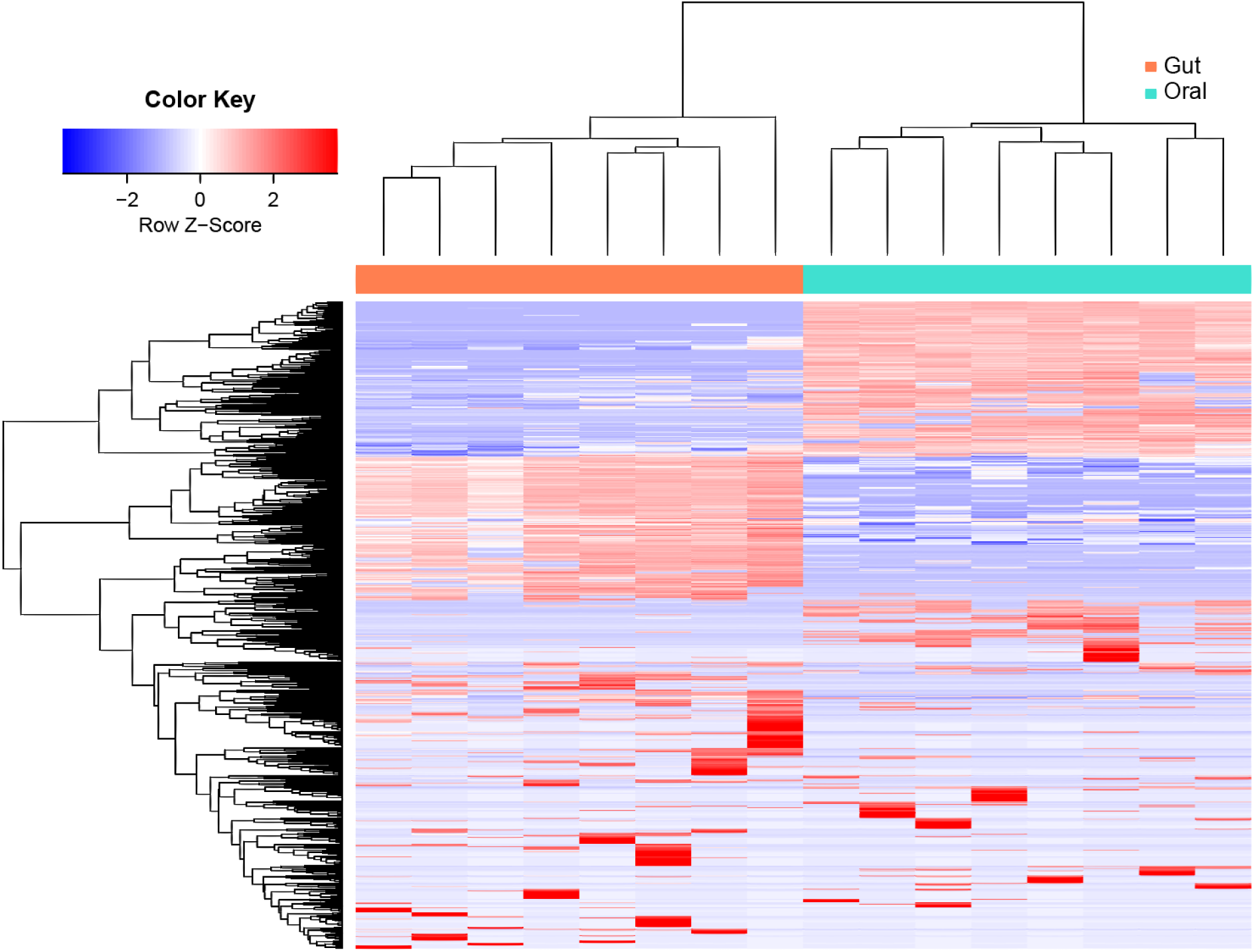
Gut and oral microbiome from the HPFS cohort.

**Figure 8:**
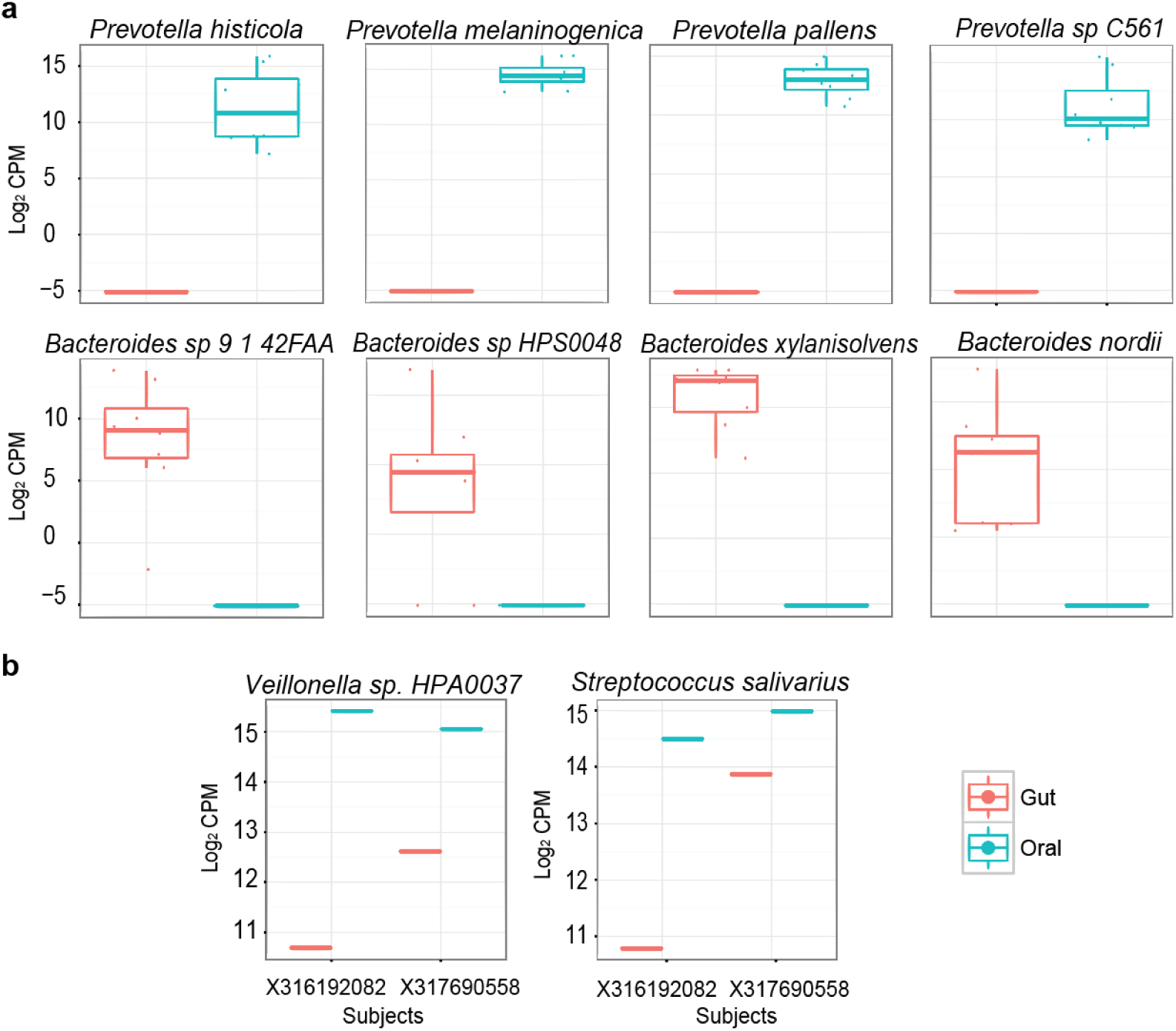
**(a)** Comparison of the species composition in gut and oral samples. The largest distinction between the oral and the gut microbiome were from several members in the *Prevotella* and *Bacteroides* genus. **(b)** Link between the oral and the gut microbiome. Two species with > 1000 CPM were found in the oral and the gut samples from the same individual (of at least two individuals): *Veillonella sp. HPA0037 and Streptococcus salivarius.*

## Discussion

The classification of homologous sequences from closely related species can be challenging in shotgun metagenomics sequencing analysis because this can lead to many misidentifications of species. In addition, samples deriving from a host may contain many contaminating host DNA, thus complicating the data analysis [15]. In recent years, many metagenomic reads classification tools have been developed with the rising use of shogun metagenomics sequencing. While most of the studies have been focusing on bacterial communities, there is also a rising use of shotgun sequencing in viral community setting [16]. In this study, we tested different pipelines with existing metagenomics and RNA-seq data analysis tools on two different microbial communities, one that mimics samples deriving from human-associated habitat, and another one that consists of only microbial reads.

In the pipelines involving the use of kallisto for quantification, we first filter out human reads using BWA-MEM. The reason why we did not use kallisto for the human filtration step was because the index could not be built from the hg19 genome due to memory issue, as kallisto is specifically designed for transcriptome. With the same set of marker genes as the reference database, kallisto was shown to outperform MetaPhlAn2 in identifying the correct species. We introduced this marker genes filtration approach in our pipeline as a species detection step to reduce false positives. We showed that without this step, many species that are not truly present were detected. However, the drawback of this pipeline is the runtime. Kallisto is a fast pseudoaligner for transcriptome [12], but it takes a much longer time for metagenome sequence classification when compared to Kraken, an ultrafast metagenomics analysis tool. However, kallisto outperformed Kraken in terms of quantification accuracy. Overall, D1 (with human filtration) and D2 pipelines demonstrated the most optimal combinations of the performance parameters that we tested.

To further test the pipelines, we applied the D2 method on iMESSi simulated DNA sequencing reads. Due to the absence of reference genomes to half of the reads present in the simulated sample, the sensitivity was not ideal. This can mimic the problem in analyzing real dataset in which there is still a problem in classifying reads extracted from unknown species. In addition, many species under the same genus were misidentified, including the *Bacillus* genus, the *Thermus* genus and the *Burkholderia* genus. This suggests that some of the markers were not truly unique to the species even after filtering out quasi-markers from MetaPhlAn2.

We also tested the D1 pipeline on real metagenomics sequence reads from the HPFS cohort (eight healthy males at the Boston area). We showed that the highest distinction between the gut and the oral samples came from members of two genus, *Bacteroides* and *Prevotella*. Several members in the *Bacteroides* genus was found in the gut but not in the oral samples and vice versa for the *Prevotella* genus. This is in agreement with previous result indicating that *Bacteroides* species in the gut microbiome is associated with an animal-protein based modern western diet whereas *Prevotella* species in the gut is associated with a carbohydrate-based rural diet [17]. We also found several *Prevotella* species that were found only in the oral site but not in the gut samples. This is consistent with the result in which *Prevotella* genus was previously shown to significantly associate with the tongue dorsum [18].

## Methods

### Generation of simulated DNA sequencing data

We simulated DNA sequencing paired-end reads to represent two different microbial communities: (i) microbial reads extracted from human host (consist of human reads and ~ 5% of microbial reads) and (ii) reads that consist of only microbial reads. To generate these reads, we used wgsim by Heng Li (https://github.com/lh3/wgsim) with hg19 genome and 315 microbial genomes (292 species: 74 bacteriophages, 69 viruses and 149 bacteria) from the NCBI RefSeq database [19]. We used the default options in wgsim except for a rate of mutation of 0.003, for the generation of microbial reads. For the simulation that contains microbial reads from human host, we generated five sets of them progressively, with approximately a total of 100 million reads (94,999,998 human reads + 5,000,001 microbial reads), 30 million reads (28,500,000 human reads + 1,499,999 microbial reads), 10 million reads (9,500,000 human reads + 499,993 microbial reads), 3 million reads (2,849,998 human reads + 149,987 microbial reads) and 1 million reads (949,997 human reads + 49,997 microbial reads). For the simulation that contains only microbial reads, we also generated five sets of reads progressively, which contain approximately 100 million reads (99,999,999), 30 million reads (30,000,001), 10 million reads (10,000,003), 3 million reads (2,999,998) and 1 million reads (999,994).

### IMESS_Illumina simulated Illumina100

In addition to our simulated sequencing reads, we also evaluated the D2 approach on the dataset that was used by Schaeffer et al. to test kallisto in a metagenomics setting [13]. This dataset was simulated using iMESS_Illumina and contains 100 bacterial genomes [20].

### Quantification of microbial reads

We compared MetaPhlAn2, kallisto/0.42.2 and kraken/0.10.5-beta for their performance in microbial reads quantification. For the simulated microbial sample of human origin, we first filtered out reads mapping to hg19 genome [21], with the exception when using MetaPhlAn2 for classification. No prior filtration is required for MetaPhlAn2 due to the use of microbial species-specific markers database. For the approach involving Kraken (E1), we directly used Kraken to filter out human reads with the options, --quick, --min-hits 5 --unclassified-out, with hg19 genome as the reference genome. Next, we ran Kraken on the unclassified reads output against the full microbial reference genomes database. For the approaches using kallisto, we compared three different ways (B1-D1) to classify microbial reads. In the first approach (B1), we ran BWA-MEM [22] against hg19 genome to filter out human reads. Then, we classify unmapped reads with the full microbial reference genomes database using kallisto. In the second approach (C1), we ran an additional step of kallisto on hg19 unmapped reads with species-specific markers from MetaPhlAn2 as the reference genomes. This step is introduced to reduce the FDR and it is used to detect the presence or the absence of a species, and not for quantification purpose. If a species is detected here, the estimated read count will be obtained from the earlier step that uses full microbial genomes database. In the third kallisto approach (D1), we again ran BWA-MEM against hg19 genome for human reads filtration. The unmapped reads were classified using kallisto, with species-specific markers from MetaPhlAn2 as the reference genomes to detect the presence of a species. After identifying species that were present in the sample, we proceeded to build a kallisto index on full microbial genomes only on these selected species. The quantification step was ran with this reduced database.

For the second microbial community that contains only microbial reads, we ran MetaPhlAn2, kallisto and Kraken exactly the same way as we ran them in the first microbial community found in human-associated habitat, but without the filtration step to remove human reads.

### Reference genomes

The hg19 reference genome was obtained from UCSC Genome Bioinformatics, University of California, Santa Cruz [23]. Due to the very high memory requirement by kallisto, we included only finished microbial sequences in our reference database based on available species-specific markers from MetaPhlAn2 for the comparisons between five pipelines [6]. 3508 finished microbial sequences that made up of 2616 microbial species, including phages, archaea, bacteria and viruses, were obtained from the NCBI RefSeq database [19] and used in kallisto and Kraken pipelines. For pipelines that involve the use of markers database, we included only markers that are found in the 3508 finished microbial sequences. Quasi-markers and the set of markers that were excluded by MetaPhlAn2 were not included in our marker database. A quasi-marker was identified as having at least one external genome where it maps to. The same set of marker genes were used in our MetaPhlAn2 pipeline. For the analysis of dataset from iMESS_Illumina simulated Illumina100 [20], we used all species-specific marker genes available at the species level from MetaPhlAn2 as our first step of analysis. These markers are from 109985 microbial reference sequences (4868 bacterial, viral and fungal species). The subsequent step of building a reduced database (Figure 9 D1 and Figure 10 D2) uses sequences from 109985 fasta entries.

**Figure 9:**
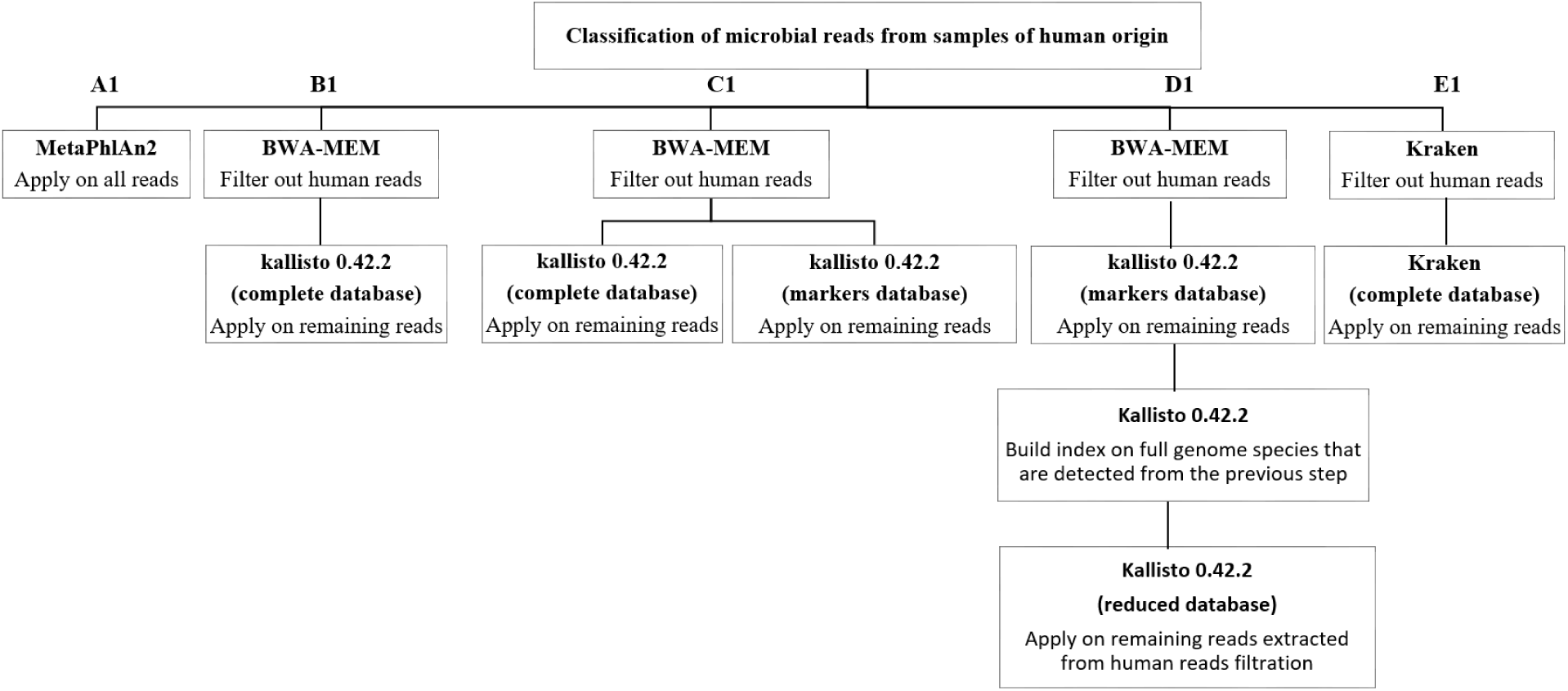
Flowchart of analysis pipelines for samples extracted from human-associated habitat. Five pipelines (A1-E1) were evaluated and compared for their overall performances.

**Figure 10:**
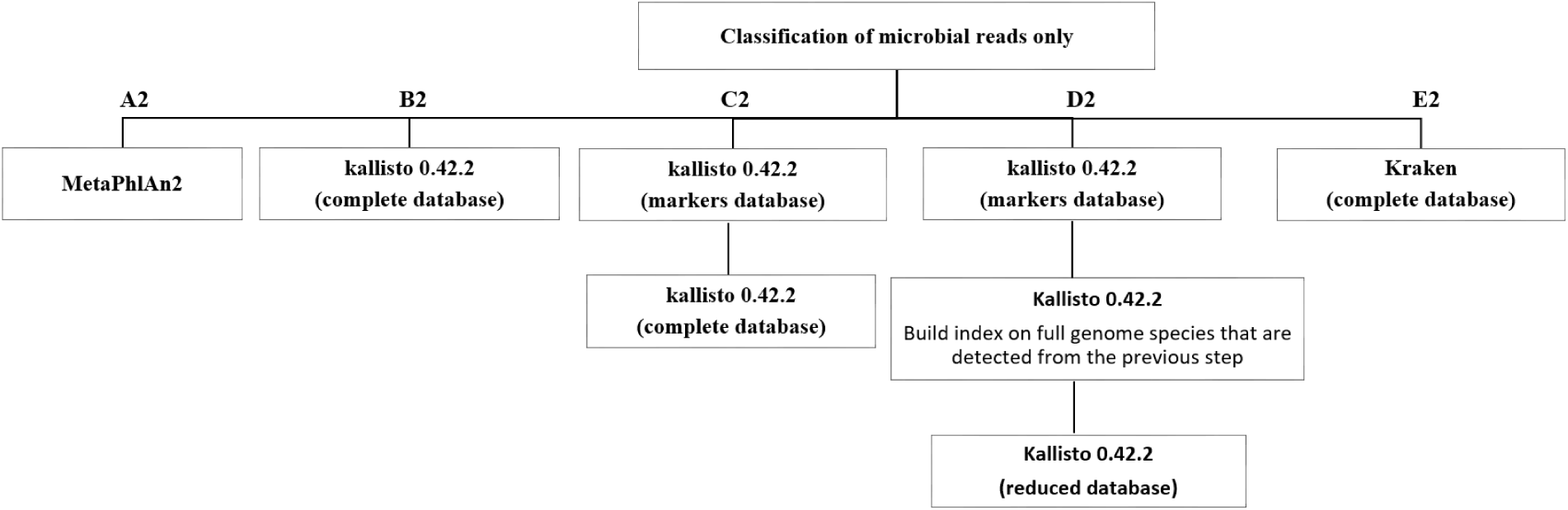
Flowchart of analysis pipelines for samples that consist of only microbial reads. Five pipelines (A2-E2) were evaluated and compared for their overall performances.

### Stool and saliva sequence data from the HPFS cohort

The stool and saliva whole genome shotgun sequencing data from eight healthy HPFS male participants were obtained from the Sequence Read Archive (accession number: PRJNA188481) [14].

### Quality Control

To extract high quality reads, we filtered iMESSi-generated reads and the DNA sequencing reads from the HPFS cohort using prinseq v0.20.4 [24] with options -min_len 40, -trim_qual_left 10, -trim_qual_right 10, -min_qual_mean 18.

### Evaluation of performance

We evaluated the five different classification approaches for their performances in terms of (i) false discovery rate (total number of species detected that were not in the dataset, out of the total number of species detected), (ii) sensitivity (total number of species detected, out of the total number of species truly present), (iii) rate of false negative (number of species that were in the dataset but not detected, out of the total number of species truly present) (iv) accuracy (correlation between estimated counts and ‘ground truth’ that is represented by RMSE) (v) memory requirement, and (vi) runtime.

### Statistical analysis

For the analysis of metagenomic data from the HPFS cohort, we used estimated counts from kallisto to calculate log_2_CPM using the EdgeR package [25]. The heatmap was generated with log_2_CPM.

